# White-Matter BOLD Mediates Time-Varying Cortico-Cortical Functional Connectivity

**DOI:** 10.64898/2025.12.04.691728

**Authors:** Lyuan Xu, Yurui Gao, Muwei Li, Zhongliang Zu, Kurt G. Schilling, Adam W. Anderson, Zhaohua Ding, John C. Gore

**Affiliations:** Vanderbilt University Institute of Imaging Science, Vanderbilt University Medical Center, Nashville, TN USA; Department of Electrical Engineering and Renewable Energy, Oregon Institute of Technology, Klamath Falls, OR USA; Department of Biomedical Engineering, Vanderbilt University, Nashville, TN USA; Department of Radiology and Radiological Sciences, Vanderbilt University, Nashville, TN USA; Department of Electrical and Computer Engineering, Vanderbilt University, Nashville, TN USA

## Abstract

Functional connectivity (FC) measurements in the brain using BOLD fMRI have largely focused on correlations between gray-matter (GM) BOLD signals, ignoring the possible relevance of white-matter (WM) in delineating neural networks. We first employed a tract-informed mediation framework to test whether WM pathways contribute indirect, bundle-specific effects to cortico-cortical coupling. Using resting-state fMRI data from the Human Connectome Project, we quantified pathway-level mediation effects across GM-WM-GM units and found that WM BOLD signals significantly mediated GM-GM associations, with clear tract-dependent heterogeneity. Building upon this, we examined how dynamic features of WM BOLD signals relate to time-varying intra-cortical FC. We found that temporal variability of WM signals, rather than their mean amplitudes, more closely tracked fluctuations in GM-GM dynamic connectivity, with correlations reaching approximately 0.4 across specific tracts. Together, these results indicate that WM BOLD signals are related to how cortical regions interact, underscoring their integral role in large-scale brain network organization.

## Introduction

Functional connectivity (FC) measurements in the brain using BOLD fMRI have largely focused on correlations between gray-matter (GM) BOLD signals, ignoring the possible relevance of white-matter (WM) in delineating neural networks. Yet converging evidence indicates that white matter (WM) BOLD fluctuations are robustly measurable and exhibit distinct patterns of activity and inter-regional correlations (1, 2), supporting the view that BOLD signals in WM are related to neural activity within the brain (3, 4). However, cortical FC is itself not stationary, and time-varying or dynamic FC has been widely reported across tasks and rest (5, 6). Moreover, although the engagement of WM in GM networks has been demonstrated (7), the contributions of WM to measured GM FC have not been quantified. We therefore aimed to assess how temporal features of WM BOLD signals are related to dynamic variations in cortico-cortical coupling, and the degree to which WM pathways modulate GM-GM interactions.

Two gaps in our current knowledge motivate the present work. First, in conventional analyses, correlations are typically assessed between cortical regions only, while the contribution of WM explicitly is ignored (8). Recent work by Zu et al. represents one first attempt to avoid this over-simplification (9). Alternatively, a tract-informed statistical mediation framework, which tests whether the functional association between two cortical regions contains an indirect component transmitted through a connecting WM bundle, may provide a compact, pathway-level summary of WM contributions. Second, conventional analyses have related BOLD time series in WM to those of GM regions, but conceptually it may be more appropriate to consider whether WM signals reflect the instantaneous connectivity between communicating GM regions rather than their activity. Different descriptors of WM signals capture different physiological and statistical properties. A running average of the mean WM signal may reflect baseline GM signal amplitudes or their correlations, leaving a key ambiguity in how WM BOLD should be summarized when probing its relationship to cortical connectivity.

We conducted a systematic investigation to characterize how WM BOLD dynamics relate to time-varying cortical FC, and whether WM pathways mediate cortico-cortical coupling at a tract level. Specifically, we first implemented a mediation framework to assess whether BOLD activity in WM bundles modulates functional associations between connected cortical regions. We then compared dynamic characteristics of WM BOLD signals, comparing running averages of their mean versus their variance, to determine which more closely aligns with fluctuations in GM connectivity over time. By integrating dynamic and mediation perspectives, this work provides a pathway-level view of how WM contributes to cortical FC.

Our findings suggest that WM pathways exhibit significant mediating effects on connectivity that are bundle specific, while the temporal variability of WM BOLD signals - not their mean amplitudes - better track dynamic GM connectivity. Together, these results offer additional evidence that WM BOLD signals are related to how cortical regions interact, underscoring their integral role in large-scale brain network organization.

## Results

We first investigated the mediating roles of WM BOLD signals in the functional connectivity between GM regions. Figure 1 illustrates average bundle-specific mediation effects from a simple first-order mediation framework in which the FC between two GM regions is decomposed into a direct component (*c*^′^) and an indirect component transmitted through a WM bundle. In this linear formulation, the total effect is *c* = *c*′+*a*⋅*b*, where *a* indexes the association between GM1 and WM, and *b* indexes the association between WM and GM2 while controlling GM1. The product *a*⋅*b* quantifies the indirect effect magnitude of functional association between two GM regions that is mediated by the corresponding WM BOLD signals. The resulting patterns are clearly bundle dependent: some tracts exhibit stronger and more distributed indirect effects, whereas others show weaker or more focal influences, broadly consistent with the roles within known large-scale networks. Although preliminary and not causal, the results provide a compact, tract-specific summary of WM contributions to cortico-cortical coupling.

**Figure 1.**
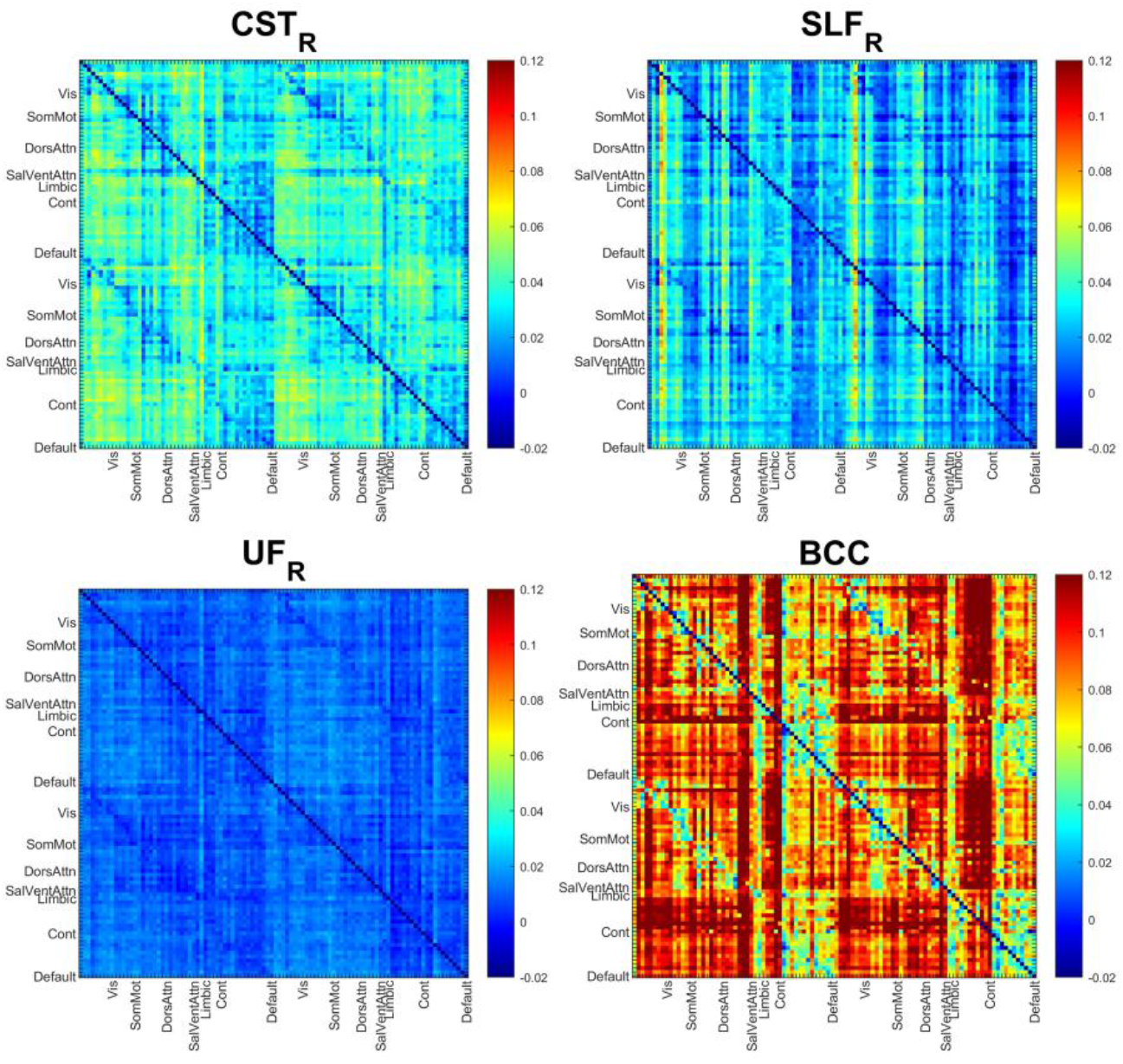
Averaged bundle-specific mediation effects of WM BOLD signals on GM-GM functional connectivity for four different WM bundles across all subjects.

We then examined the correlations between the dynamic GM-GM FC and dynamic features derived from WM BOLD signals within each sliding window, including both the windowed variation and mean, as shown in Figure 2. The corresponding error bars are also shown in the figure. The results show that, for all corresponding GM-WM-GM units, the mean correlation coefficients (CC) between WM BOLD variation and dynamic GM-GM FC are consistently higher than that between mean WM BOLD signals and dynamic FC, as indicated by the orange and blue points. For the latter, most values remain near zero. Furthermore, for each GM-WM-GM unit, we concatenated the time-domain curves obtained from sliding windows across all subjects and performed the same dynamic analysis without averaging. The resulting correlations between WM BOLD variation and dynamic GM-GM FC, shown as red dots, increased further compared with the averaged values (orange dots), with some CC up to approximately 0.4. The findings may suggest that the low-frequency power of fluctuations in WM BOLD, rather than signal amplitude, is an important factor driving cortical FC.

**Figure 2.**
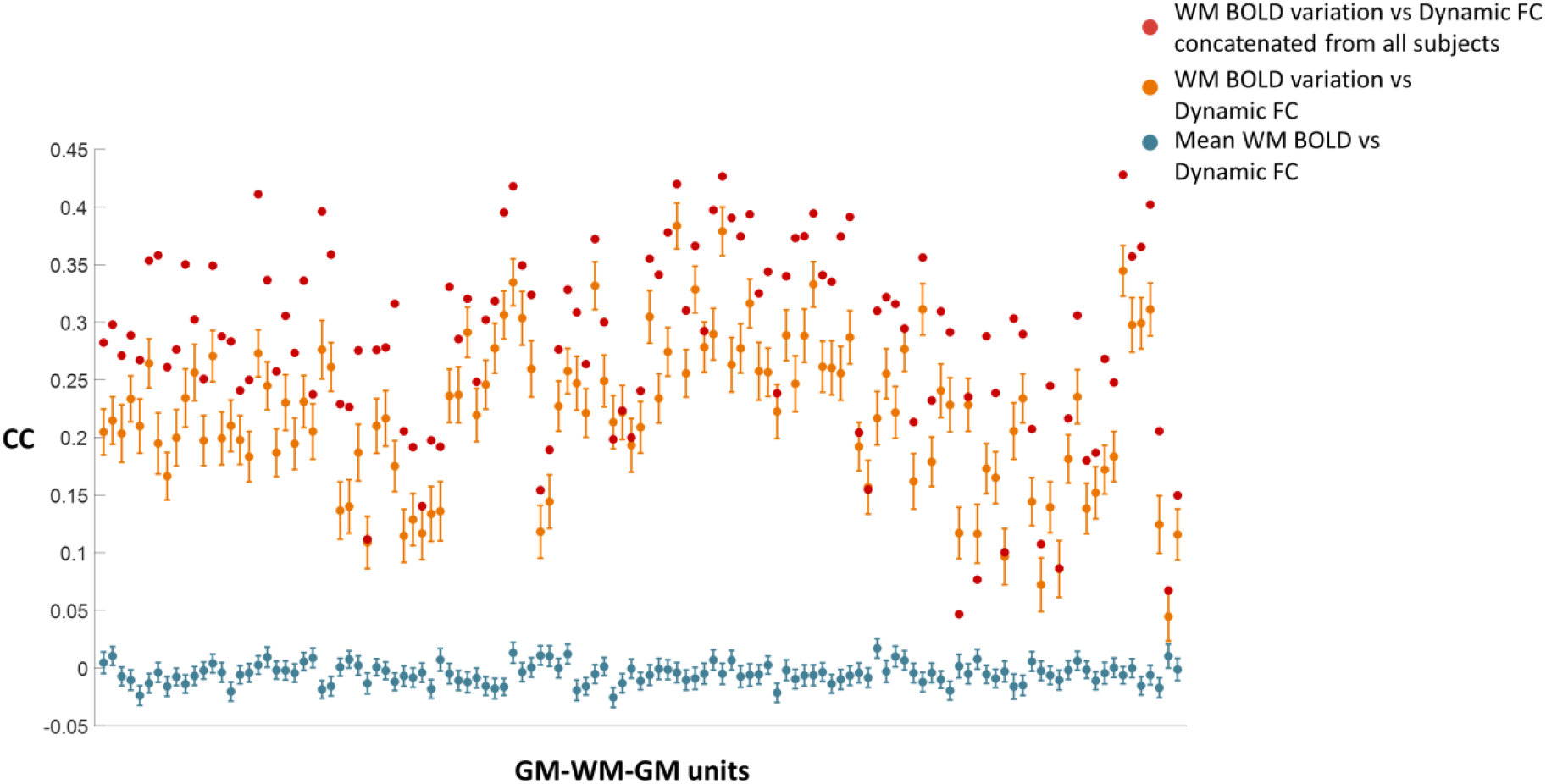
Correlations between WM BOLD dynamics and dynamic GM-GM FC. The averaged correlation coefficients (CC) between dynamic GM-GM FC and mean WM BOLD signal, across all subjects for different GM-WM-GM units, are close to zero. By contrast, the CCs between dynamic FC and the variance of WM BOLD signals are significantly larger than zero and in some tracts are approximately 0.4.

To further determine the statistical significance of the observed results, we employed a permutation test for validation. Across all GM-WM-GM units concatenated over participants, the mean correlation between WM BOLD-variance and the dynamic FC of the corresponding GM pair was 0.284 (as shown by the red line in Fig. 3). To assess whether this association could arise from random WM–GM pairing, we conducted a permutation test that preserved each dynamic GM-GM FC series while randomly reassigning the intervening WM bundle, recomputed the correlation for each triad, averaged across triads, and repeated this procedure to form a null distribution. The observed mean correlation lay well to the right of this distribution, exceeding its 95th percentile (as shown by the yellow dashed line), indicating a non-random, tract-specific coupling between WM signal variability and GM dynamic connectivity.

**Figure 3.**
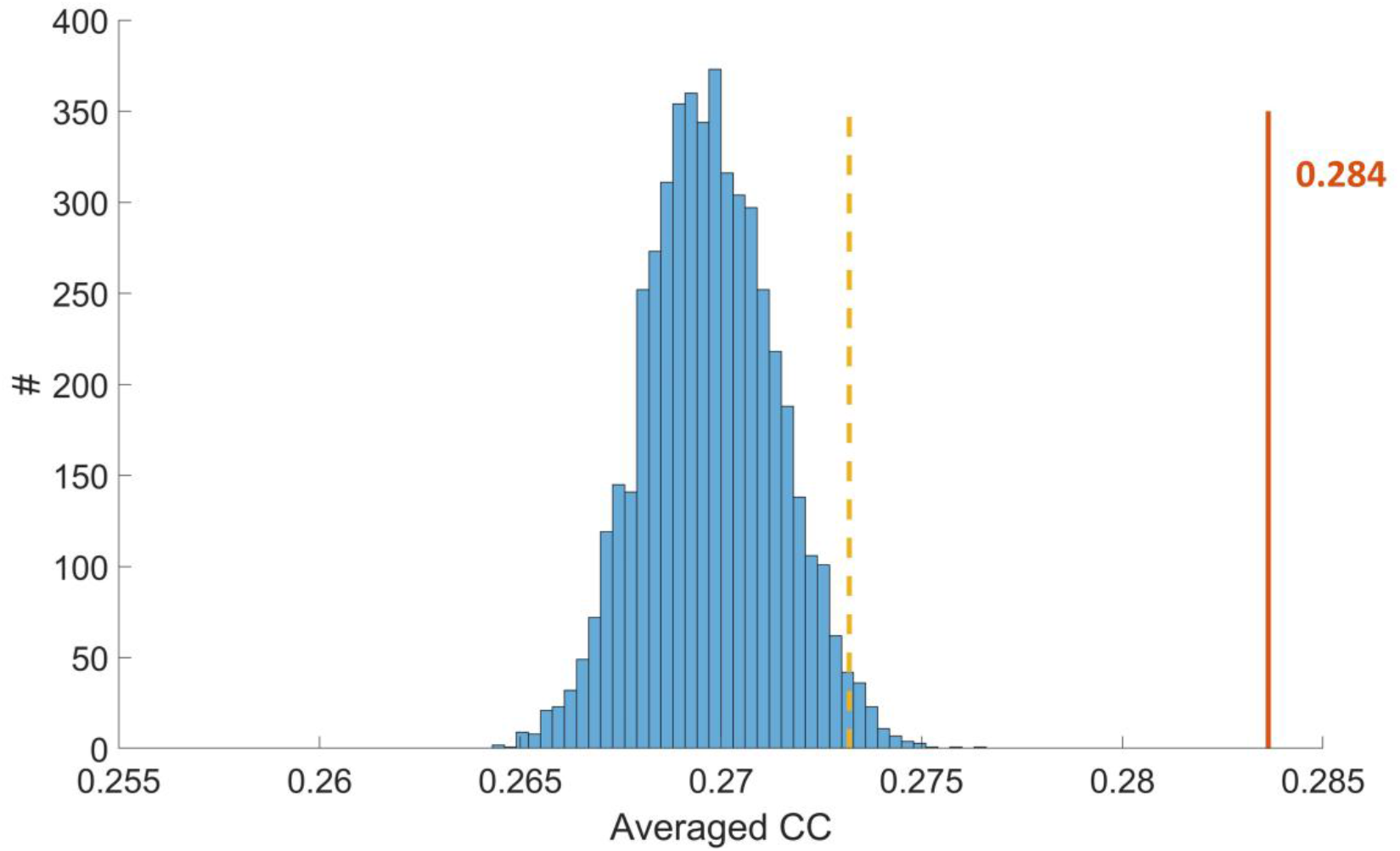
Averaged CC between WM BOLD variation and dynamic FC concatenated from all subjects across all GM-WM-GM units (0.284, labeled as orange) and the distribution of averaged CC obtained from 5000 random permutations of WM bundles (labeled as blue). The position corresponding to the 95th percentile is marked with a yellow dashed line.

**Figure 4.**
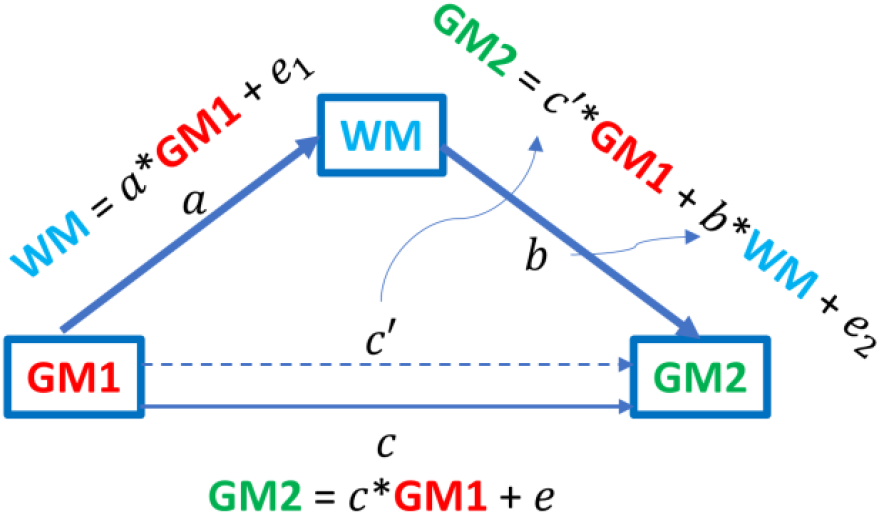
Mediation analysis model illustrating the modulatory role of WM in GM-GM functional connectivity. In this framework, GM1 and GM2 represent the independent and dependent variables, respectively, and WM serves as the mediator. Path coefficients (*a, b, c, c*^′^) represent the effects of GM1 on WM, WM on GM2, the total effect of GM1 on GM2, and the direct effect of GM1 after accounting for WM. The mediation effect (*a* · *b* = *c* − *c*^′^) quantifies the portion of the GM-GM functional relationship that can be attributed to WM-mediated modulation.

## Discussion

In this study we examined whether specific tracts contribute an indirect, pathway-dependent contribution to GM-GM coupling and how WM BOLD activity aligns with time-varying cortico-cortical FC. A tract-informed mediation framework yields bundle-dependent indirect effects that compactly summarize how WM mediates GM-GM associations beyond the direct effect. The mediation term provides a compact summary of pathway-specific, indirect effects attributable to the WM bundle linking two cortical regions. We observe clear bundle dependence in these indirect components - some tracts exhibit stronger and more distributed mediation than others - suggesting WM pathway contributes to GM-GM associations. Also, temporal variability of WM BOLD (windowed variance) tracks within-subject fluctuations in GM-GM FC more consistently than the windowed mean amplitude. Across GM-WM-GM units, correlations between dynamic GM-GM FC and the variance of WM BOLD were reliably above zero, whereas correlations with the mean WM signal were near zero. When time series were concatenated across participants to emphasize temporal resolution, several tracts approached correlation magnitudes of ∼0.4. These patterns indicate that fluctuation magnitude, rather than baseline level, is the WM descriptor that most faithfully aligns with changes in cortical coupling.

The modulatory role of WM in neural activities of brain networks is a continuing area of research, with accumulating evidence indicating that WM BOLD fluctuations participate actively in modulating cortical network dynamics. As shown by Ding et al. (2025), WM BOLD fluctuations reflect spontaneous cortical network activity, and their modulation contributions vary substantially across different network regions. These spatial variations are closely associated with cortical structural properties, such as local myelin content (10). In addition, some studies have also demonstrated the crucial role of WM within functional brain networks, rather than merely serving as a mediator or representing a nuisance component (7). In this study, we employed a mediation analysis framework to examine the mediating role of WM in cortical connectivity, and found heterogeneous effects across different WM regions, which may suggest WM participates actively in modifying functional network organization and this mediation role may also be shaped by its underlying functional specialization.

Additionally, we emphasize the dynamic characteristics of WM, particularly the windowed variance, which showed a remarkable concordance with GM-GM functional connectivity. This further underscores the indispensable and critical modulatory role of WM within functional brain networks. It is worth noting that it was the windowed variance of WM BOLD signal, rather than its mean value, that exhibited a stronger correlation with the cortical dynamic FC, which may be attributed to several factors. Esfahlani et al. demonstrated that FC is not uniform but is often driven by a limited number of high-amplitude co-fluctuation events (11). This directly supports the notion that the energy envelope of WM, rather than its mean activity, underlies the variation of cortical functional coupling. In addition, it has been shown that BOLD signals are more closely related to the power spectrum of local field potentials (LFPs) (12). Therefore, the windowed variance of WM BOLD signals - which reflects their underlying low-frequency power - may more effectively capture the covariation with cortical BOLD dynamic FC.

This study has several limitations. First, the mediation effects observed here do not necessarily imply causal relationships; factors such as temporal delays or external confounds should be carefully considered. In addition, the windowing approach may exhibit varying sensitivity across different frequency-related components, and the development of more adaptive joint time-frequency domain analytical methods could be beneficial. Nevertheless, the present work links WM dynamic fluctuations to cortical dynamic coupling and, through a tract-level mediation framework, delineates the pathway-specific contributions of WM. Collectively, these findings further clarify the functional role of WM dynamics in large-scale network organization.

## Materials and methods

### Data and preprocessing

MRI datasets used in this study were obtained from the Human Connectome Project (HCP) database, comprising 120 healthy young adults (50% female, 50% male). Following the approach of Xu et al. (13), to minimize potential seasonal effects, the number of subjects in each seasonal subgroup was kept identical. Functional MRI data from the HCP repository had undergone minimal preprocessing (see details in (14)). Additional preprocessing steps included the regression of nuisance variables related to head motion, cardiac, and respiratory signals using the PhysIO toolbox (15). Finally, the time series were band-pass filtered to retain frequencies between 0.01 and 0.1 Hz and normalized voxel-wise to unit variance.

### Regions of interest

The basic FC analyses in this study were performed at a regional level. For GM, functional regions of interest (ROI) were defined using Schaefer 100 atlas (16). Meanwhile, for WM, in the mediation analysis, we applied the JHU ICBM-DTI-81 WM atlas (17) to parcellate WM fiber bundles. The preprocessed fMRI signals were averaged across voxels within each GM region and WM bundle, yielding region-averaged signals. For the dynamic analysis, we defined a GM-WM-GM unit as a structural triplet comprising two GM parcels and the WM bundle connecting them. Specifically, over 260,000 WM streamlines were extracted from the HCP1065 population-averaged tractography atlas (18) and reassigned to specific WM bundles based on their connected GM pairs. The detailed procedure for streamline assignment is described in our previous work (19).

### Mediation analysis of WM BOLD signals on GM-GM connectivity

To quantitatively evaluate the contribution of WM in modulating FC between GM regions, we performed a mediation analysis based on a standard first-order mediation framework. In conventional FC analysis, the correlation between BOLD signals of two GM regions, denoted as GM1 and GM2, can be represented as a direct association (GM1 → GM2). Mediation analysis introduces an intermediate variable, forming a pathway GM1 → WM → GM2, in which GM1 BOLD serves as the independent variable, GM2 BOLD as the dependent variable, and WM BOLD as the mediator linking the two. In this study, we hypothesized that the FC between GM1 and GM2 is partially mediated by specific WM bundles. For each GM1-WM-GM2 triplet, we constructed linear regression models to estimate the interrelations among their BOLD signals. Specifically, the WM signal was first regressed on GM1 signal, yielding coefficient *a* which represents the effect of GM1 on WM; then GM2 signal was regressed simultaneously on both WM and GM1 signals to obtain coefficients *b* and *c*^′^, representing the effects of WM and the direct effect of GM1, respectively; finally, the total effect *c* of GM1 on GM2 was estimated without including WM as a mediator. The mediation effect, defined as the product *a* × *b* = *c* − *c*^′^, reflects the portion of effects between GM2 and GM1 that can be attributed to WM-mediated modulation, independent of the direct influence of GM1. This measure thus provides a meaningful and quantitative indicator of the extent to which WM pathways mediate or support GM-GM FC.

### Dynamic analysis

For each GM-WM-GM unit, dynamic FC between the two GM ROIs was computed using a sliding-window approach. The window length was set to 60 s. The sliding window step size was set to 1 TR. Within each window, we also derived dynamic measures of WM BOLD activity, including the mean BOLD magnitude and temporal variation across the window. For the correlation analysis, the temporal correlations between the dynamic FC of GM pairs and the corresponding dynamic WM measures were calculated, as was shown in Figure 5. We also performed permutation tests. For each GM-WM-GM unit, the dynamic FC between GM pairs was kept fixed, while the corresponding WM bundle was randomly reassigned to another WM bundle. The correlation coefficients (CC) between dynamic GM-GM FC and WM BOLD variation (concatenated across all subjects) were recalculated for each permutation. This procedure was repeated 5000 times to generate a null distribution of mean CC values across all units.

**Figure 5.**
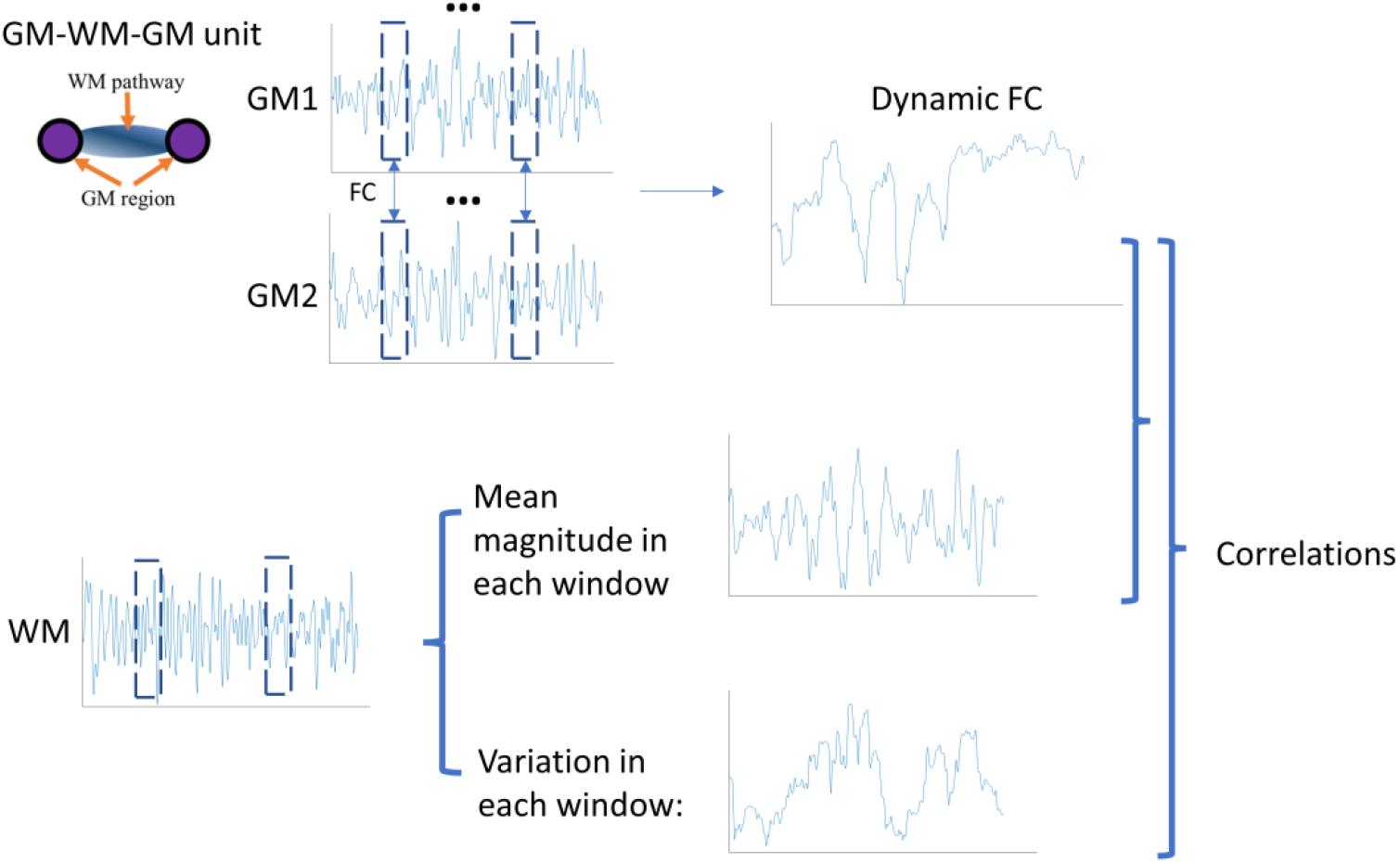
Schematic of dynamic analysis within each GM–WM–GM unit. Dynamic FC between two GM ROIs was estimated using a sliding-window approach. Within each temporal window, WM BOLD mean magnitude and temporal variance were extracted, and correlations were computed between the GM dynamic FC and corresponding WM dynamic measures.

## Acknowledgement

This work was supported by NIH grants R01NS113832 (Gore), RF1MH123201 (Gore and Landman), R21AG083915 (Gao), R01NS129855 (Ding) and K01EB032898 (Schilling).

## Ethics Approval

The studies involving humans were approved by Institutional Review Board of Washington University School of Medicine. The studies were conducted in accordance with the local legislation and institutional requirements. The participants provided informed consent to participate in this study.

